# Cross-institutional HER2 assessment via a computer-aided system using federated learning and stain composition augmentation

**DOI:** 10.1101/2024.01.17.576160

**Authors:** Chia-Hung Yang, Yung-An Chen, Shao-Yu Chang, Yu-Han Hsieh, Yu-Ling Hung, Yi-Wen Lin, Yi-Hsuan Lee, Ching-Hung Lin, Yu-Chieh Lin, Yen-Shen Lu, Yen-Yin Lin

**Affiliations:** JelloX Biotech Inc.; No. 66-5, Shengyi 5th Rd., Zhubei City, Hsinchu County 302041, Taiwan; National Taiwan University Hospital; No.7, Chung Shan S. Rd., Zhongzheng Dist., Taipei City 100225, Taiwan

**Keywords:** Human epidermal growth factor receptor 2 (HER2), federated learning, stain variation, augmentation, stain normalization

## Abstract

The rapid advancement of precision medicine and personalized healthcare has heightened the demand for accurate diagnostic tests. These tests are crucial for administering novel treatments like targeted therapy. To ensure the widespread availability of accurate diagnostics with consistent standards, the integration of computer-aided systems has become essential. Specifically, computer-aided systems that assess biomarker expression have thrusted through the widespread application of deep learning for medical imaging. However, the generalizability of deep learning models has usually diminished significantly when being confronted with data collected from different sources, especially for histological imaging in digital pathology. It has therefore been challenging to effectively develop and employ a computer-aided system across multiple medical institutions. In this study, a biomarker computer-aided framework was proposed to overcome such challenges. This framework incorporated a new approach to augment the composition of histological staining, which enhanced the performance of federated learning models. A HER2 assessment system was developed following the proposed framework, and it was evaluated on a clinical dataset from National Taiwan University Hospital and a public dataset coordinated by the University of Warwick. This assessment system showed an accuracy exceeding 90% for both institutions, whose generalizability outperformed a baseline system developed solely through the clinical dataset by 30%. Compared to previous works where data across different institutions were mixed during model training, the HER2 assessment system achieved a similar performance while it was developed with guaranteed patient privacy via federated learning.

## Introduction

Recent progression in precision oncology, including immunotherapies and antibody-drug conjugates, has showed a great potential to prolong the survival of cancer patients. To aim with the proper treatment strategies, these advances were usually accompanied by careful inspection of the patient information, such as high-throughput sequencing and companion diagnostics. Such inspection has largely benefitted from artificial intelligence (AI) and deep learning (DL) models, which provided quantitative details of biomarker expressions [1–5].

In metastatic breast cancer treatment, human epidermal growth factor receptor 2 (HER2) has played an important role in protein-targeted therapies. The HER2 target monoclonal antibody drug trastuzumab (market name: Herceptin) has become a mature first-line treatment for HER2-positive patients, reducing the risk of progression and death significantly in comparison to chemotherapies alone [6–9]. Assessing the HER2 expression involved immunohistochemical (IHC) staining and an *in situ* hybridization test if the former was deemed equivocal [10]. However, recent study has suggested that clinical HER2 interpretation could suffered from low concordance among pathologists, while with the assistance of several AI algorithms, it became more likely for the pathologists to reach an agreement [11–12]. The urgent need of AI adaption in the HER2 expression analysis was thereof apparent.

Several challenges remained when developing an AI system for HER2 expression assessment. First, DL models integrated in such a system must be data-driven, but medical images have often been scarce resources for an individual institution. More critically, policies of patient privacy [13] has prohibited data to be shared across institutions, so DL models could not be trained in a conventional manner. Fortunately, the emerging federated learning (FL) [14] has empowered medical institutions to collectively develop DL models while maintaining patient privacy. Specifically, in an FL strategy, multiple institutions iteratively trained a model with their own data in parallel, and then they communicated the model parameters with each other between iterations. This technology has demonstrated significant improvements in generalizability of various DL models for medical imaging [15–17].

Second, despite the success of FL applications on computed tomography and magnetic resonance imaging [18–20], the performance of DL models specialized for histological images could be impeded due to the imaging variation across different institutions. Histological images, e.g., IHC staining required for the HER2 expression assessment, consisted of color channels that convolved pixel-wise intensities of hematoxylin and diaminobenzidine (DAB). The color composition of these stains was heavily influenced by laboratory conditions under which the specimens had been prepared. For instance, Figure 1 demonstrated the histological images collected from various sources; although the observed stain-color variation has been well-aware by pathologists, it was hardly perceived by an DL model that strictly took the color channels as input. As a result, the trained model might become less performant and leaned toward only part of its data sources.

**Fig. 1.**
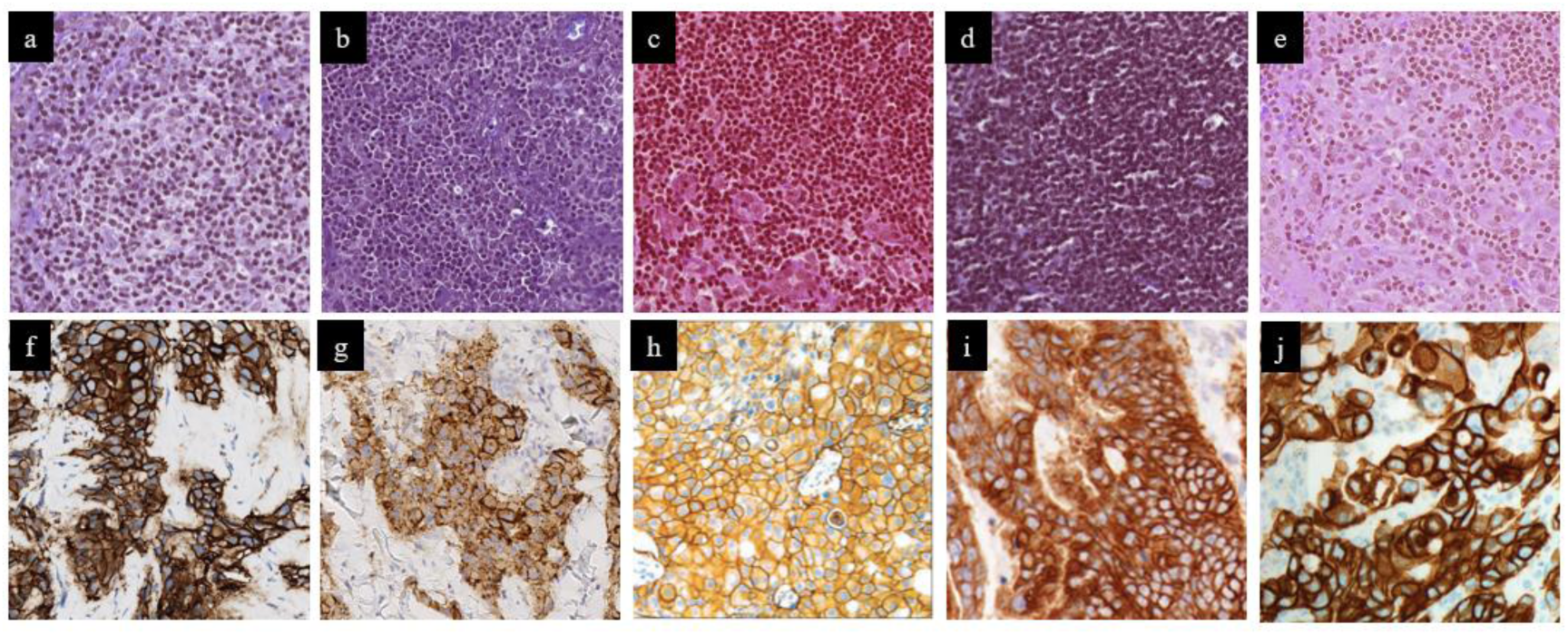
Stain variation across institutions. The top row is hematoxylin and eosin (H&E) ROIs from the public Camelyon17 dataset. (a) Canisius Wilhelmina Hospital. (b) Rijnstate Hospital. (c) University Medical Center Utrecht. (d) Radboud University Medical Center Nijmegen. (e) Laboratory of Pathology East Netherlands. The second row is HER2 IHC ROIs. (f) National Taiwan University Hospital. (g) University of Warwick. (h) HercepTestTM Interpretation Manual. (i) Bond OracleTM HER2 IHC System Interpretation Guide. (j) Interpretation Guide for VENTANA anti-HER2/neu (4B5).

In the presented work, a computer-aided HER2 assessment system was proposed, which incorporated a few techniques to overcome the obstacles of inter-institutional model development: (a) FL with a median aggregation strategy [21] was applied to a tumor segmentation model trained on both clinical and publicly available IHC slides, (b) a new technique term the stain composition augmentation was designed to amend the generalizability of the tumor segmentation during FL training process, improving the segmentation accuracy on the inferior dataset from 85% to 93%, and (c) various computer vision algorithms were combined into a HER2 membrane staining classifier, where a stain normalization procedure [22] was performed for inter-institutional viability. The HER2 assessment system was evaluated on the clinical and the public dataset, both achieving an accuracy greater than 90%. In addition, the proposed AI system showed a great flexibility of interpreting HER2 expression, which shed light on its potential contribution to the rising disputation of a new HER2 ultra-low subtype.

## Materials and methods

### Datasets

This study included breast cancer IHC slides with HER2 information that were collected from two distinct sources. First, the National Taiwan University Hospital (NTUH) dataset, with approval from the Institutional Review Board (202004032RSB), included clinical whole-slide images from 154 specimens. After formalin fixation and paraffin embedding, these collected samples were sectioned and stained with IHC using the Ventana HER2 4B5 assay on a Bench-Mark XT autostainer (Ventana Medical Systems). The digital image of each section was scanned using a VENTANA DP 200 slide scanner at 20X magnification, which resulted in 154 IHC slides. The clinical HER2 interpretation of these slides were also available.

Second, the Warwick dataset [23] consisted of 50 cases in the Her2 Scoring Contest held in conjunction with the Pathological Society of Great Britain and Ireland meeting in Nottingham (June 2016), whose IHC slides and the corresponding HER2 scores were publicly available. These slides were scanned with a Hamamatsu NanoZoomer C9600 at 40X magnification.

The IHC slides in the NTUH and the Warwick datasets were divided into the training, validation, and testing sets that consisted of 58, 15, 81 slides and 24, 6, 20 slides respectively. To develop the tumor segmentation model and the membrane staining classifier that formed the HER2 assessment system, regions of interest (ROIs) at 20X magnification were cropped from the tissue regions within the IHC slides. Specifically, for the tumor segmentation model, the tumor regions in the IHC slides were annotated by four well-trained scientist (YHH, YLH, YWL, FTC) and reviewed by an experienced pathologist (YHL). 4267 and 2810 ROIs of size 1280-by-1280 were selected from the NTUH and the Warwick training/validation dataset respectively. For the computer vision algorithm of membrane staining classification, 112 and 140 ROIs of size 320-by-320 were cropped from the NTUH/Warwick training dataset respectively. No annotations were created for these ROIs, but they showed roughly comparable occupations of negative/faint/weak/strong membrane staining judged by the well-trained scientist.

### Stain composition augmentation

Here, an augmentation technique was proposed to simulate a variety of histological images based on a source image, which could thereof enlarge the coverage of data distribution during model training. Particularly, histological images differ from regular RGB images such that, in an ideal scenario, their pixels were only colored by two stains, e.g., hematoxylin and DAB for IHC images. Both the difference of staining intensity and the color composition of stains could lead to variability of histological images across institutions. The proposed technique, termed the stain composition (SC) augmentation, followed these two scenarios to generate augmented images that are more aligned with the distribution of histological data compared to existing augmentations which focused on RGB images in general.

Given an input IHC image, the SC augmentation began with calibrating its hematoxylin and DAB channels through color deconvolution [22] (Fig. 2a). The color deconvolution algorithm decomposed the IHC image *X* into its color basis of the stains *S* and the corresponding staining intensity *A*:

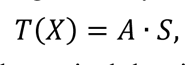

where *T* transformed the image to the optical density space [24], and *S* consisted of unit-length row vectors representing the color composition of hematoxylin and DAB, along with an orthogonal background vector.

**Fig. 2.**
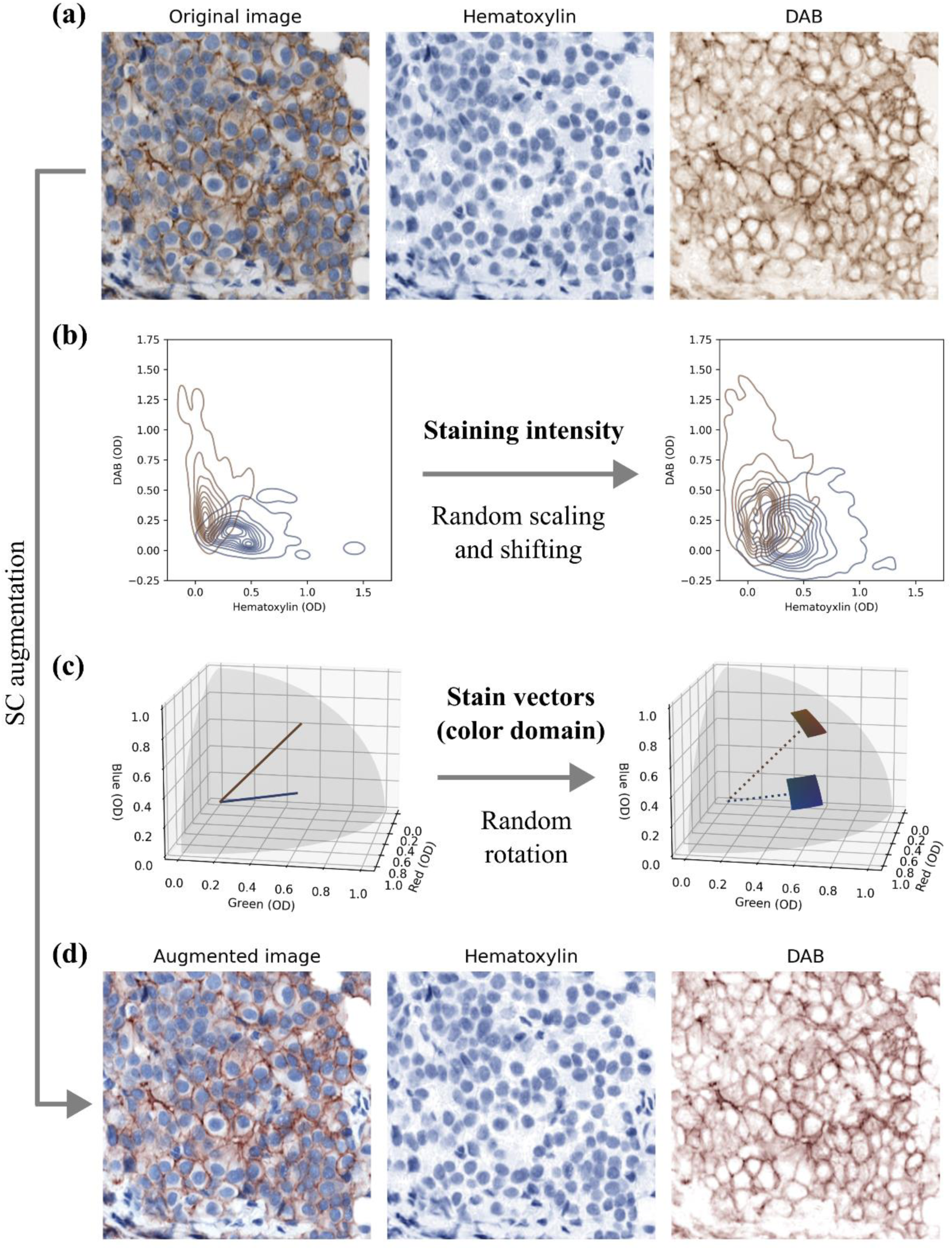
Procedure of the SC augmentation. An IHC image was decomposed into its hematoxylin/DAB channels using color deconvolution (a). The intensity and the color composition of staining were randomly varied (b-c), which led to an augmented image analogous to an IHC image collected from a different source (d).

Next, the intensity *A_ij_* of every stain *j* and pixel *i* was scaled by a random factor *α_j_* and shifted by a random offset *β_j_*:

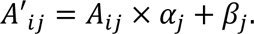

The variation of staining intensity of the IHC image was thus increased (Fig. 2b). Furthermore, the color composition *S_j_* for each stain *j* was perturbed. Supposed *(1, φ(S_j_), θ(S_j_))* to be the spherical coordinates of *S_j_*, it was rotated by random angular shifts:

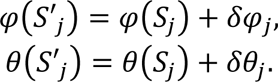

The angles *φ(S’_j_)* and *θ(S’_j_)* were then clipped between 0 and *π/2*, so the perturbed color composition *S_j_* remained in the valid optical density space. This perturbation randomly varied the color domain of hematoxylin and DAB in the proximity of the original IHC image (Fig. 2c). All the scaling *α_j_*, shifting *β_j_*, and rotation parameters *δφ_j_* and *δθ_j_* were randomly sampled from uniform distributions with some predetermined ranges. Combined, the augmented image became (Fig. 2d):

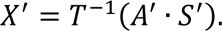

The proposed SC augmentation was further evaluated with the public Camelyon17 challenge dataset [25], which consisted of H&E slides collected from five medical centers, and it was compared to six existing augmentation/normalization techniques. In Supplementary Figure S2, the SC augmentation was showed more capable of bridging the gap of distinct distributions of histological images, which also led to an improved stability and generalizability of a metastasis classification model trained using federated learning.

### Computer-aided HER2 assessment system

The presented computer-aided HER2 assessment system integrated various techniques to overcome the challenge of inter-institutional data variation and patient privacy protection, including federated learning, stain composition augmentation and stain normalization, as illustrated in Fig. 3. Specifically, the entire system consisted of two major modules: a deep learning model of tumor segmentation and a computer vision algorithm for nuclei detection and membrane staining classification.

**Fig. 3.**
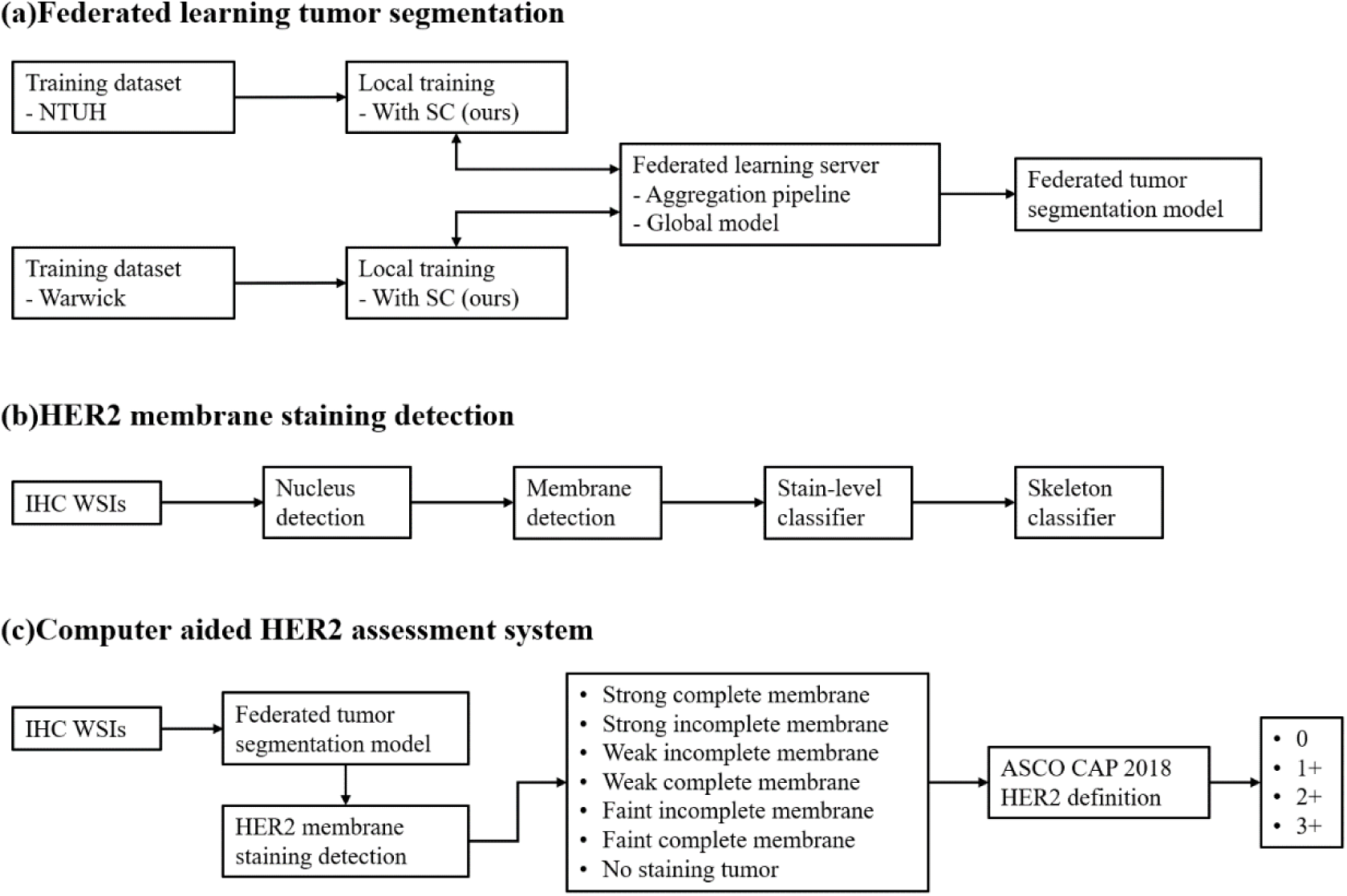
Overview of the HER2 assessment workflow. The development process was divided into three parts: (a) Federated deep learning training to establish a tumor segmentation model. (b) Computer vision analysis of membrane staining to classify each cell membrane. (c) The HER2 WSIs scoring process involved integrating the results from parts (a) and (b) for quantitative statistics and determining the final scores in accordance with the ASCO CAP 2018 HER2 Guideline.

### Federated learning tumor segmentation

(Fig. 3a) A convolutional deep learning model was developed to identify regions of tumor cells. This model was trained through federated learning platform between the NTUH and the Warwick datasets. In particular, the training procedure took place over 20 epochs, where, after every 1 epoch of parameters updates within each dataset locally, the median aggregation strategy [21] was employed to combine the model parameters learned from the NTUH and the Warwick data distribution. In addition, the proposed SC augmentation was applied throughout the federated learning process, so the model featured an increased stability during federated training and an improved generalizability across institutions.

Our tumor segmentation model followed a revised version [26] of the HRNetV2 architecture [27]. The model was trained with the mean squared error objective, an Adam optimizer, and a batch size of 20 images. An exponential decay learning rate scheduler is used as well, starting at 0.001 and decaying by a ratio of 0.96. The training images were down-sampled to 5X magnification and were randomly cropped to a size of 160-by-160 pixels. Basic augmentation techniques were applied, including horizontal and vertical flipping, ninety-degree rotations, brightness/contrast adjustment [28], and elastic deformation [29]. Furthermore, during model training, the SC augmentation was sampled from uniform distributions ranging between −0.1 and 0.1 for parameters *α*, *β*, *δφ*, and *δθ*, except that *δθ* of hematoxylin ranged between −0.2 and 0.2 instead. The federated training was conducted via the Intel Open Federated Learning Toolkit [30].

When predicting the tumor region of an IHC image, the image was down-sampled to 5X magnification, and the trained tumor segmentation model was applied to every 160-by-160 tile. Next, the tile-level segmentations were spliced into the corresponding locations in the original image, and the combined segmentation was subsequentially postprocessed with Otsu thresholding [31], median blurring, and up-sampling back to the image’s original magnification level.

### HER2 membrane staining detection

(Fig. 3b) First, nuclei in an IHC image were detected. A scalar feature was computed for each pixel via combining the intensity of hue, saturation, value, and the pixelwise variance of the RGB intensity. The segmentation mask of nuclei was generated by Otsu thresholding [31] on this scalar feature. Moreover, the segmentation of nuclei was divided into the segmentation mask for each nucleus through two consecutive splitting procedures, each of which involved a connected component analysis, local peak detection, and the watershed algorithm [32–33]. The uncovered nucleus components were further filtered by their size and roundness.

Second, with the nuclei detected in an IHC image, the segmentation mask of each nucleus was first dilated to obtain its membrane region. The degree of dilation was dynamically calculated based on approximating the membranes as hexagonal packing of circular objects.

Third, the membrane region of each nucleus was classified by its staining level Pixels within the membrane region were transformed into their hematoxylin-DAB representation through color deconvolution [22]. Subsequently, they were classified into the negative, faint, weak, and strong levels based on three predetermined thresholds on the DAB channel. We then classified each membrane as the strongest staining level of its pixels that had sufficient presence in the membrane region.

Forth, the membrane regions were further classified by their completeness of staining. For each membrane, a skeletonization algorithm [34] was applied to the pixels corresponding to the staining level of the membrane region and the pixels whose staining were weaker by one level. The skeleton contours were then processed by subsequential closing operations, and a membrane was categorized as complete if the skeleton contour was closed and enclosed a significant area of the membrane; otherwise, it was deemed incomplete.

The parameters of the computer vision algorithm of nuclei detection and membrane staining classification was tuned by visually inspecting the prediction on 112 ROIs and 7 WSIs selected from the NTUH training dataset. The details of these tuned parameters were documented in Supplementary Table S1.

Besides, when utilizing these computer vision algorithms on images of the Warwick dataset, a stain normalization procedure is applied beforehand to enhance generalizability of the algorithms. Specifically, the color domain of hematoxylin and DAB was estimated by color deconvolution [22] for tuning ROIs in the NTUH and the Warwick dataset respectively. The Warwick images were transformed into the NTUH color domain by combining the NTUH stain-color composition with the stain intensity extracted from the Warwick color domain (which was scaled with a predetermined factor for each stain).

### Computer-aided HER2 assessment system

(Fig. 3c) The inter-institutional, computer-aided HER2 assessment system was design as such: The tumor regions in an IHC slide were first uncovered by the tumor segmentation model. Next, using the computer vision algorithms, each nucleus in the tumor regions was detected and its membrane region was classified as negative, faint-incomplete, faint-complete, weak-incomplete, weak-complete, strong-incomplete, or strong-complete. Based on the distribution of membrane staining in the tumor regions, an IHC slide was interpreted as 0, 1+, 2+, or 3+ according to the ASCO CAP 2018 guideline [10].

## Results

The tumor segmentation model was separately evaluated with annotated IHC slides in the NTUH and the Warwick dataset (Fig. 4). When the model had been trained with federated learning but without the proposed SC augmentation, it achieved a decent accuracy of 94.82% on the Warwick dataset; nevertheless, the model demonstrated a compromised accuracy of 85.39% on the NTUH dataset. On the other hand, combining federated learning and the SC augmentation, the tumor segmentation model reached an accuracy of 96.82% and 92.68% for the Warwick and the NTUH dataset respectively. The SC augmentation advanced the overall accuracy of the tumor segmentation model from 89.29% to 94.39%.

**Fig. 4.**
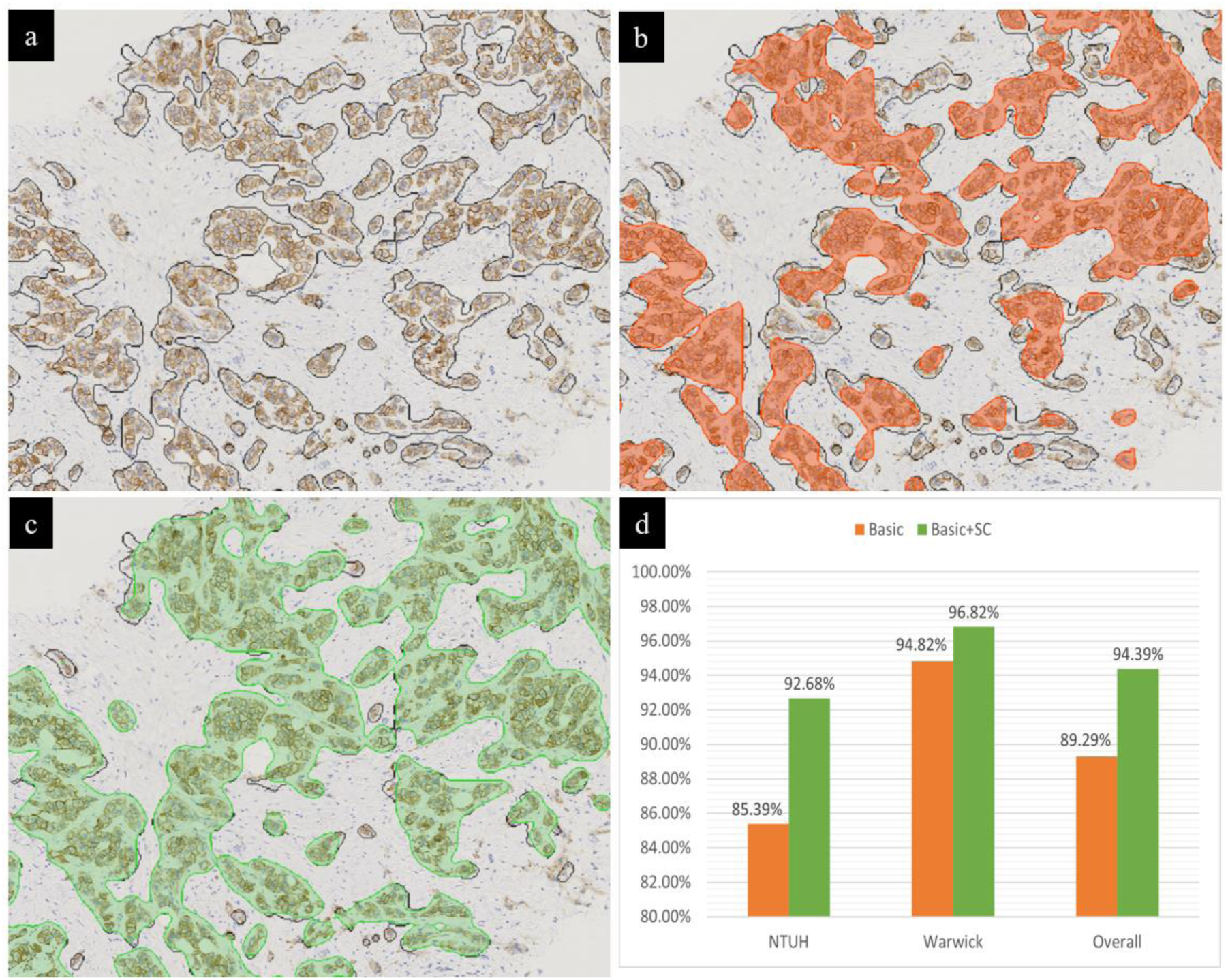
Performance of tumor segmentation model trained with FL between NTUH/Warwick images. (a) An example ROI selected from the Warwick dataset along with its tumor region annotation. (b) Predicted tumor regions (orange) when the SC augmentation was absent in FL. (c) Predicted tumor regions (green) combining FL and the SC augmentation. (d) Accuracy of tumor segmentation with/without the SC augmentation.

Figure 4 depicted the predicted tumor regions of an example IHC slide. It was apparent that, when the SC augmentation had been absent from the federated training, the prediction neglected many tumor cells. Incorporating the SC augmentation mitigated the imbalance of color domains of the IHC images during model training. As a result, the tumor segmentation model became generalizable across different institutions.

The presented HER2 assessment system fulfilled a good performance across the two institutions by combining the generalizable tumor segmentation model and the membrane staining classification algorithm along with stain normalization preprocessing. Figure 5 showed the confusion matrix and various evaluation metrics, where our HER2 assessment system achieved an accuracy of 90% (45/50) and 85% (17/20) on the NTUH and the Warwick dataset respectively. Should the prediction be grouped into negative (0/1+), equivocal (2+), and positive (3+) as in current clinical practices, the assessment system even achieved an accuracy of 94% (47/50) and 90% (18/20) respectively. The cross-institution generalizability of our assessment system was also manifested when visualizing its prediction of example IHC slides (Fig. 6); regardless of the stain variation between the NTUH and the Warwick IHC slide, the membrane staining classification algorithm could identify the level of HER staining and its staining completeness.

**Fig. 5.**
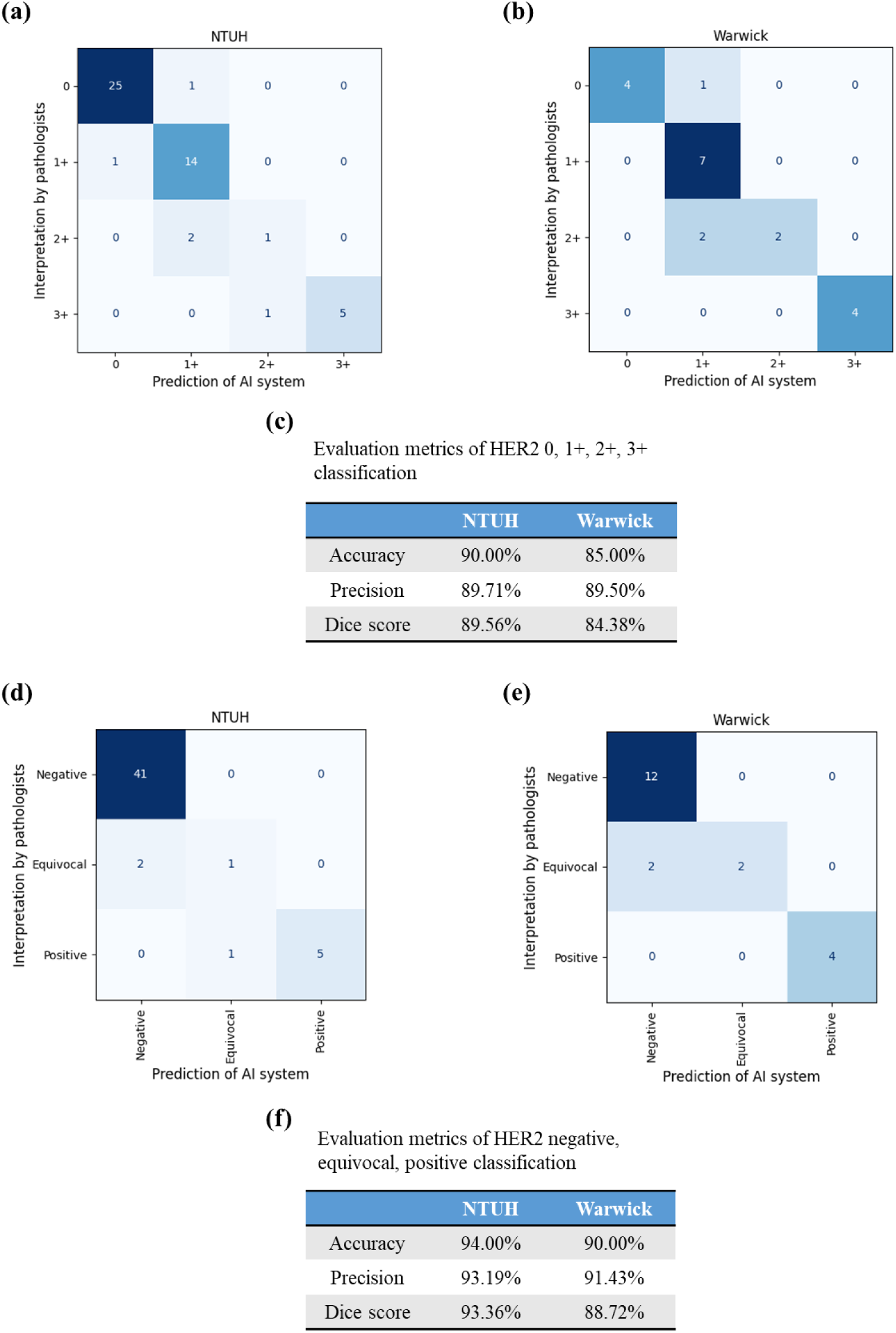
Confusion matrix and evaluation metrics of the HER2 assessment system. The prediction of the HER2 assessment system was compared to the pathologists’ interpretation. Two sorts of classification were presented: (a-c) classification among the HER2 0, 1+, 2+, and 3+ grading, and (d-e) classification among HER2 negative, equivocal, and positive.

**Fig. 6.**
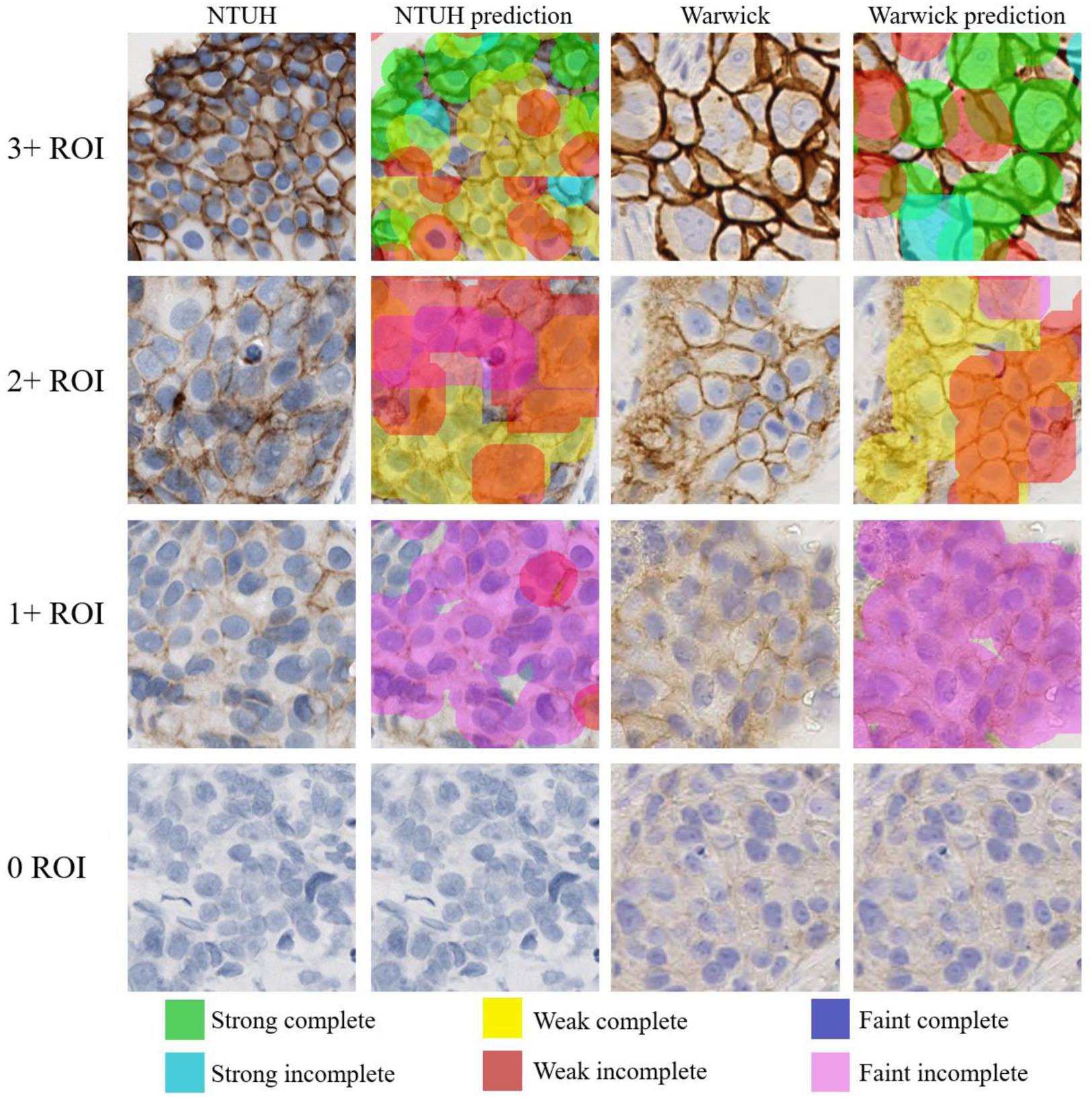
Membrane staining classification in example tumor regions. Eight ROIs were selected from the NTUH and the Warwick dataset, and the prediction of membrane staining generated from the HER2 assessment system was presented with different colors according to their classification.

Furthermore, the proposed HER2 assessment system was compared with a baseline system which was developed solely with IHC slides in the NTUH dataset (see Supplementary materials). This baseline system was also separately evaluated with the NTUH and the Warwick dataset, which achieved an accuracy of 86% and 55% respectively (Fig. S4). Such a comparison suggested the techniques of federated learning, the SC augmentation, and the stain normalization preprocessing could greatly increase the system’s generalizability. By incorporating data collected from other institutions, the accuracy of HER2 assessment could be slightly amended by 4% on the NTUH dataset. More critically, these techniques regulated the stain variation across different institutions and led to an assessment system applicable to IHC slides of various sources, such as the Warwick dataset for which the accuracy was improved from 55% to 85%, by a 30% increment.

## Discussion

Inter-institutional generalizability has long been a challenge for AI systems in medical imaging applications, especially for quantitative applications in precision medicine. In this study, a variety of techniques were integrated into a biomarker computer-aided framework to overcome the challenge of inter-institutional generalizability, which led to a HER2 assessment system applicable to both a clinical and a publicly available dataset. Using federated training, a DL tumor segmentation model was jointly optimized by images of different sources. The FL procedure was further amended by a newly proposed augmentation approach, where the hematoxylin/DAB staining in the IHC images was independently varied so the model could accommodate the stain variation between institutions. Furthermore, for a computer vision algorithm of HER2 membrane staining classification, the HER2 assessment system leveraged stain normalization so the classifier could be adapted to images of various sources. Such a normalization procedure remained critical since many biomarker AI systems to date [12,35–36] involved traditional algorithms that lacked a way to be jointly optimized by mutually inaccessible datasets.

We shall address that the presented work merely showcased a preliminary success to develop an AI system of HER2 expression assessment that could be generalized to different medical institutions, and a few limitations should be noted. First, the proposed methodology strongly depended on the extent to which the color domain and the intensity of stains could be extracted from IHC slides. Recent advances of color deconvolution and the subsequent stain normalization [37] can be incorporated to further improved the models’ generalizability, such that the algorithm could tolerate slight differences in the specimen preparation protocol and the selection of antibody/scanner would be less restricted. Second, model design refinement could make the HER2 assessment system more aligned with pathologists’ interpretation. For instance, the current tumor segmentation model was not capable of dissecting ductal carcinoma *in situ* (DCIS) from invasive tumor regions, where the former was recommended to be excluded in HER2 interpretation [10], and another DL model detecting the DCIS regions could be integrated. Third, in addition to evaluating the HER2 assessment system by its concordance with the pathologists’ interpretation, future investigation would be required to assess how this computer-aided system could increase the concordance between pathologists of different institutions [12].

Despite its aforementioned limitations, the presented HER2 assessment system was able to achieve a comparable performance to existing studies. Glass et al. [38] extensively annotated the HER2 expression of individual cancer cells in 689 slides of ductal carcinoma that were stained and digitalized across multiple laboratories, and the resultant model reached an intraclass correlation coefficient (ICC) of 0.91 (95% CI between 0.89 and 0.94) on the 172 testing slides. Wu et al. [12] assessed their HER2 scoring algorithm on 246 HER2 0/1+ cases collected by a single medical institution, which achieved an accuracy of 93%. In contrast to these previous works, the proposed HER2 assessment system explicitly enforced the policy of patient data privacy through federated learning, and it led to an ICC of 0.95 (95% CI between 0.92 and 0.97) and a 94% accuracy among HER2 0/1+ cases (50/53). Additionally, in alignment with the two previous works where faint membrane staining were easier to be detected by AIs than by the pathologists, our assessment system also showed sensitivity to HER2 expression such that 25% (2/8) of the discordant cases were HER2 0 slides being classified into HER2 1+ by the system.

Moreover, the substantial HER2-expression sensitivity of the AI system indicated its great potential to assist contemporary drug development routines. As clinical studies revealed promising response of the antibody-drug conjugate Trastuzumab Deruxtecan (market name: Enhertu) in HER2-low patients [39–40], it has been disputed that whether a new HER2 ultra-low subtype should be considered in HER2 interpretation [41–43]. However, the definition of this new subtype is still under investigation, which has been confronted with difficulties to precisely identify the barely perceivable HER2 expression in IHC slides [44]. The proposed AI system quantified HER2 membrane staining of various characteristics, and it was flexible to comply to the adapting definition of the HER2 ultra-low subtype (see Fig. 7), for which the lower threshold of faint membrane staining has remained unclear [45].

**Fig. 7.**
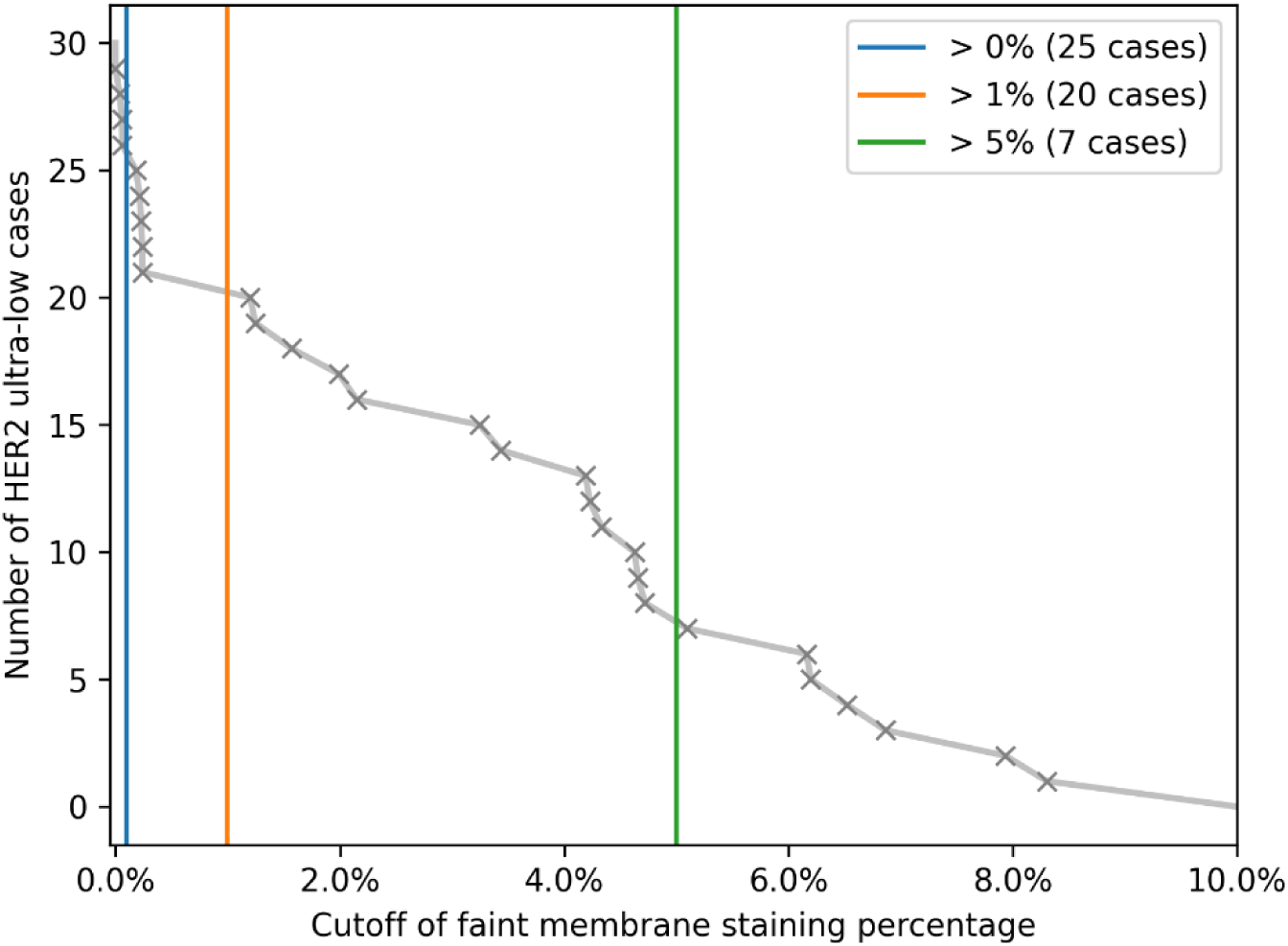
Inversed cumulative distribution of faint membrane staining in HER2 0 cases. The number of HER2 ultra-low cases found in the NTUH/Warwick dataset was computed with several hypothetical cutoffs of the percentage of faint membrane staining in the IHC slides.

Particularly, in the NTUH and the Warwick datasets, it was found that 65% (20/31) of the HER2 0 cases had showed faint membrane staining among 1-10% of the tumor cells. When developing the interpretation guideline of a new drug, the AI system could quickly estimate the outcomes of different changes, e.g., adding a new subtype, and the costly repetitive interpretation by pathologists could be thereof avoided.

## Conclusions

In this work, an artificial intelligent HER2 assessment system with generalizability across different medical institutions was proposed. The AI system leveraged federated learning to simultaneously analyze the characteristics of HER2 expression in IHC images collected from a clinical and a publicly available dataset. Furthermore, a new augmentation approach was designed to overcome the variation of staining among different institutions that could have biased the federated learning procedure. Combined, the HER2 assessment system achieved an accuracy more than 90% when being evaluated on IHC slides of both sources.

## Acknowledgement

We thank the staffs of National Taiwan University Hospital, and fellows Fang-Ting Chang (FTC) of JelloX Biotech Inc. for their technical assistance.

## Supplementary materials

### Comparison between stain composition augmentation and existing approaches

#### Existing Stain-Color Augmentations/Normalizations

We experimented several existing augmentation and normalization approaches with federated models, included deep learning normalization techniques StainGAN [1] and StainNet [2], augmentation on the hue-saturation-value (HSV) color space of images [3,4], and a basic augmentation incorporating horizontal and vertical flipping, ninety-degree rotations, brightness and contrast adjustment [5], and elastic deformation [6].

We also included two other techniques that had inspired our proposed augmentation approach. First, with known color information of the hematoxylin, eosin, and DAB staining [7], Tellez et al. [8] transformed the RGB values of pixels into their stain intensities. The magnitude of each stain was scaled by a random factor and shifted by a random bias, and the adjusted stain intensities were combined with the constant color composition of the stains to form the augmented image. This approach was named as the hematoxylin-eosin-DAB (HED) augmentation.

Second, Macenko et al. [9] suggested an unsupervised approach to find the color domain of the two major stains from a collection of histological images. The uncovered stain-color composition was distribution specific, and an image could be transformed from its source distribution to a target distribution by combining the target stain-color composition with the stain intensity calculated from the source composition. We referred to this approach as the color deconvolution (CD) normalization.

#### Datasets

We assessed augmentation/normalization techniques on H&E images collected by multiple institutions, which were released through the public Camelyon17 challenge [10]. The Camelyon17 challenge cohort consisted of 50 thoroughly annotated H&E slides of breast lymph node resections, which were collected from breast cancer patients across five medical centers (10 slides each). These centers were abbreviated as alphabets A to E, corresponding to the following institutions: Canisius Wilhelmina Hospital, Rijnstate Hospital, University Medical Center Utrecht, Radboud University Medical Center Nijmegen, and Laboratory of Pathology East Netherlands. The tumor metastases were delineated by experts. The slides were scanned 0.25 µm/pixel resolution and were down-sampled to 10X magnification. 22,074 non-background patches were extracted (consisting of 5192 positive and 16882 negative ones), which were split into training, validation, and testing sets of a 6:1:2 ratio.

As what Fig. S1 demonstrated, on the hue-value projection plane, some institutions in the Camelyon17 challenge (see Datasets) presented distinguishable distributions. After applying the SC augmentation, the distributions became more overlapping due to the magnified hue variation, which would be less achievable in the case of the HED augmentation that lacked stain decoupling and perturbations on color domains.

**Fig. S1.**
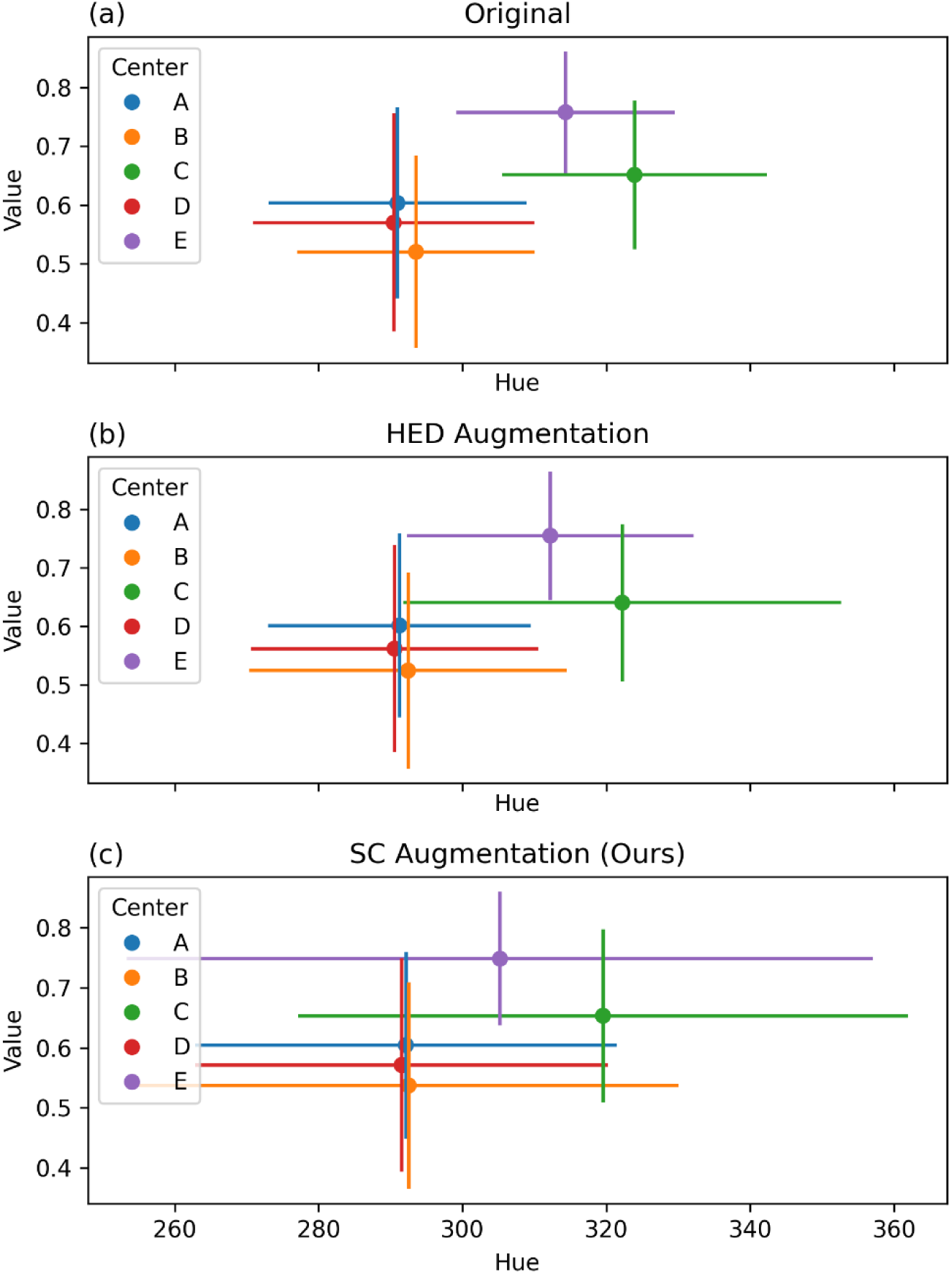
Hue-value projection of original and augmented images, where solid dots and lines represent the mean and standard deviation respectively.

#### Model Implementation

We combined models of two different tasks with augmentation/normalization approaches. For H&E images in the Camelyon17 challenge, we trained a metastasis classification model that identified whether an image patch contained metastatic tumor cells. For IHC images in the Warwick and the NTUH datasets, a tumor segmentation model was trained to detect the tumor region. Federated training of both models was conducted via the Intel Open Federated Learning Toolkit [11].

The metastasis classification model was of the EfficientNetV2S architecture [12]. The input image shape was set to 256-by-256 and the model output scores for the positive and negative classes. The model was trained with the binary cross-entropy loss and the Adam optimizer, along with an exponential decay learning rate scheduler starting at 0.001 and decaying by a ratio of 0.96 every 3,000 steps. We used a batch size of 20 and trained the model for 8,000 steps under federation of the average aggregation strategy [13].

For StainGAN and StainNet normalization, we trained the StainGAN neural network following [14] and utilized it to prepare the StainNet network following [15]. The uniform distributions, from which the basic, HSV, and HED augmentations were sampled, were parameterized the same as the experiments done by Tellez at el. [16]. The SC augmentation was sampled from uniform distributions ranging between −0.1 and 0.1 for parameters *α*, *β*, *δφ*, and *δθ*, except that *δθ* of hematoxylin ranged between −0.2 and 0.2.

#### Experiments

Here, we performed federated training across Center B and C in the Camelyon17 challenge. A metastasis classification model was trained accompanied by each of the seven augmentation/normalization techniques, and the trained models were evaluated on the testing images of internal institutions (Center B and C) and on the whole dataset of external institutions (Center A, D, and E). Due to the dramatic difference of numbers of positive and negative samples, we assessed model performance by the average precision (AP), which is the area under the precision-recall curve. Compared to the area under the receiver operating characteristic curve, the AP is a more sensitive metric in the cases of class imbalance.

Fig. S2 demonstrates the models’ performance with respect to each individual institution and to the internal/external institutions as a whole. When being evaluated on all the internal (column BC) or all the external institutions (column ADE), our proposed SC augmentation achieved the highest AP among all the approaches, scoring 0.8699 and 0.938 respectively. In addition to the superior generalizability across institutions, the SC augmentation facilitated federated training to be more stable; specifically, even for the institution where the model was least performant internally (Center B) and externally (Center D), it still achieved AP of 0.8107 and 0.9549 respectively.

**Fig. S2.**
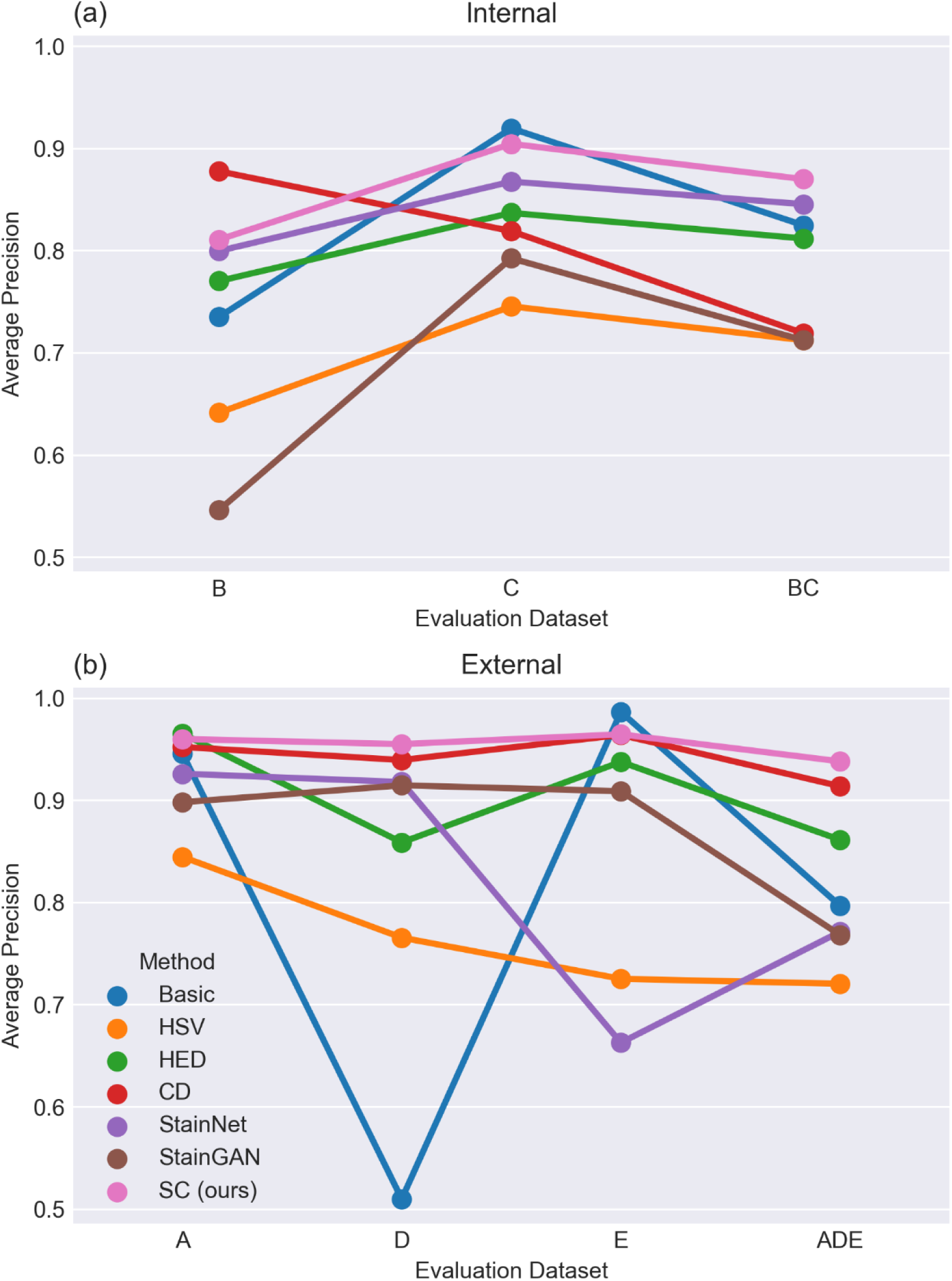
Average precision of the FL model combined with different techniques.

Other existing augmentation/normalization techniques also ameliorated federated training in our experiments, but their resultant improvements were more limited compared to the proposed SC augmentation. The basic augmentation hardly assisted the model to generalize to Center D; the HSV augmentation reduced performance variation across institutions, but its generalizability was inferior; the HED augmentation and the CD normalization led to decent model improvements on external institutions whereas the former only had internal boosts comparable to the basic augmentation and the later showed inferior overall internal performance (column BC); the StainGAN and StainNet normalization biased toward the majority and performed poorly on Center B and Center E respectively.

We further compared the performances of federated models with those under the hypothetical scenario where data could have been shared across institutions and models could have been trained with the “mixed” data. This scenario has usually been regarded as a baseline to assess the effectiveness of FL [17]. Fig. S3 shows the model AP under federation and the hypothetical mixed training respectively, when being combined with an augmentation/normalization method. The AP difference between each pair of models was displayed in text. We observed that most augmentations and normalizations had led to moderate gains of mixed training AP. More importantly, the proposed SC augmentation reduced the performance difference between federated and mixed training, which out-competed others on both internal and external evaluations.

**Fig. S3.**
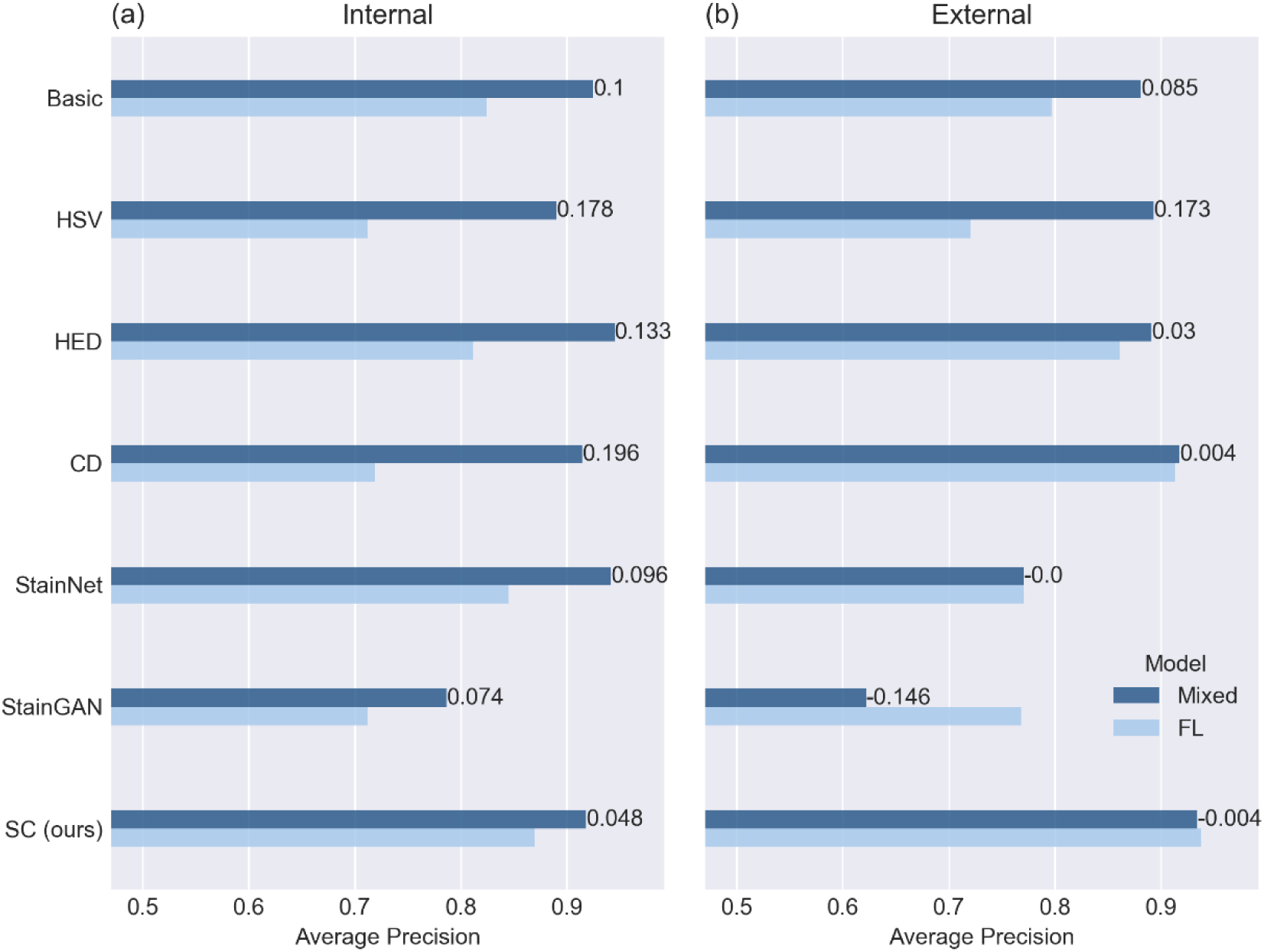
Comparison of FL model performances to the hypothetical mixed training scenario.

#### Hyperparameters of HER2 membrane staining classification

As mentioned in the Methods and materials section, the computer vision algorithm of membrane staining classification was tuned with images in the NTUH dataset. The detailed parameter values after tuning were presented in Table S1.

**Table S1.** Parameters of the computer vision algorithm for nucleus detection and membrane staining classification.

| Nucleus detection |  |
| --- | --- |
| Hue shifting | -20 |
| Minimum component size | 35 |
| Threshold of aspect ratio | 0.6 |
| Threshold of extent | 0.6 |
| Color deconvolution |  |
| Threshold of optical density | 0.15 |
| Extreme percentile for minimum/maximum | 1 |
| Scaling factor during normalization (hematoxylin) | 1.5 |
| Scaling factor during normalization (DAB) | 1 |
| Membrane staining classification – staining level |  |
| Threshold of DAB (faint staining) | 91 |
| Threshold of DAB (weak staining) | 141 |
| Threshold of DAB (strong staining) | 206 |
| Cutoff of membrane area proportion (faint staining) | 0.05 |
| Cutoff of membrane area proportion (weak staining) | 0.05 |
| Cutoff of membrane area proportion (strong staining) | 0.2 |
| Cutoff of staining pixels proportion (faint staining) | 0 |
| Cutoff of staining pixels proportion (weak staining) | 0.2 |
| Cutoff of staining pixels proportion (strong staining) | 0.3 |
| Membrane staining classification – completeness |  |
| Kernel size of closing | 3 |
| Number of closing iterations | 3 |
| Threshold of area enclosed by skeleton contour | 0.3 |

#### Baseline system of HER2 assessment

Here, a baseline HER2 assessment system was developed from image data solely in the NTUH dataset. This baseline system was then evaluated on both the NTUH and the Warwick dataset, which, in comparison to the performance of the proposed HER2 assessment system, entailed the generalizability improvement contributed by various techniques used in the proposed framework. Specifically, similar to the proposed system, the baseline system consisted of a tumor segmentation model and a membrane-staining classification algorithm as well. Nonetheless, the tumor segmentation model was simply trained by the NTUH images using the conventional approach, and the SC augmentation was not involved in model training; the membrane-staining classifier was not accompanied by any stain normalization preprocessing, and it was also optimized through the NTUH images, in analogy to the proposed system. Figure S4 showed the performance of the baseline system evaluated on the two datasets. It was found that, without the techniques of federated learning, the SC augmentation, and stain normalization, the baseline HER2 assessment system performed poorly on images collected by another medical institution.

**Fig. S4.**
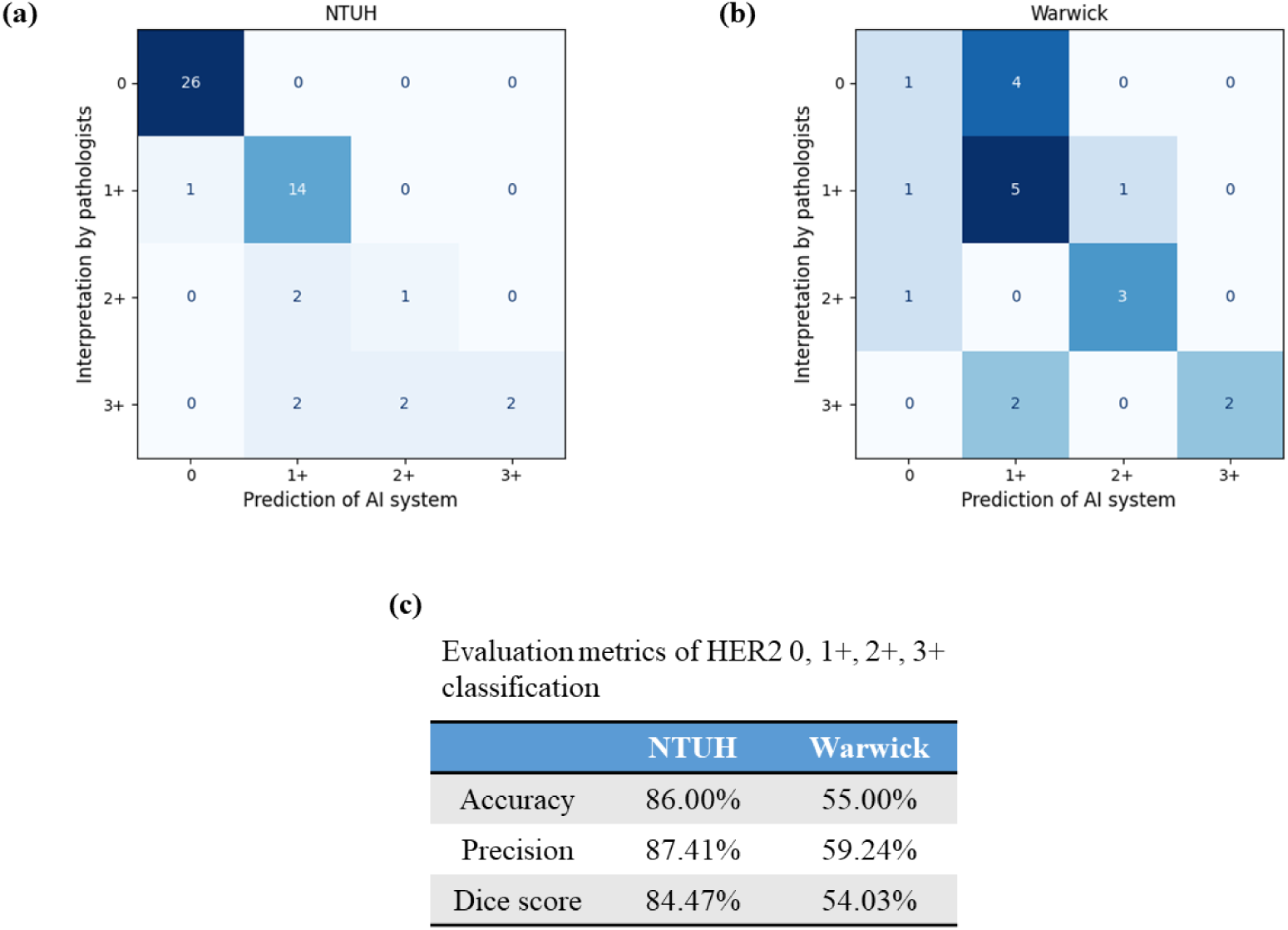
Confusion matrix and evaluation metrics of the baseline HER2 assessment system. The baseline HER2 assessment system was evaluated with the NTUH and the Warwick dataset separately, where prediction of the baseline system was compared to the pathologists’ interpretation.

